# Association of Body Index with Fecal Microbiome in Children Cohorts with Ethnic-Geographic Factor Interaction: Accurately Using a Bayesian Zero-inflated Negative Binomial Regression Model

**DOI:** 10.1101/2024.05.17.594725

**Authors:** Jian Huang, Yanzhuan Lu, Fengwei Tian, Yongqing Ni

## Abstract

The exponential growth of High-Throughput Sequencing (HTS) data on the microbial communities presents researchers with an unparalleled opportunity to delve deeper into the association of microorganisms with host phenotype. However, this growth also poses a challenge, as microbial data is complex, sparse, discrete, and prone to zero-inflation. Moreover, current methods for integrating microbiome data and other covariates are severely lacking. Hence, we propose a Bayesian zero-inflated negative binomial (ZINB) regression model that is capable of identifying differentially abundant taxa with distinct phenotypes and quantifying the effects of covariates on these taxa. Our model exhibits excellent performance when tested on simulated data. Upon successfully applying our model to a real multi-ethnic cohort study, we discovered that the prevailing understanding of microbial count data from previous research was overly dogmatic, because only a subset of taxa demonstrated zero inflation in real data. Moreover, we have discovered that dispersion parameters significantly influence the accuracy of model results, and increasing sample size can alleviate this issue. In all, we have presented an innovative integrative Bayesian regression model and a comprehensive pipeline for conducting a multi-ethnic cohort study of children, which facilitates bacterial differential abundance analysis and quantification of microbiome-covariate effects. This approach can be applied to general microbiome studies.

**IMPORTANCE:** Microbiome are closely associated with physical indicators of the body, such as height, weight, age and BMI, which can be used as measures of human health. How to accurately identify which taxa in the microbiome are closely related to indicators of physical development is valuable as microbial markers of local child growth. Complex biological systems can be effectively modeled with ZINB model which is a Bayesian Generalized Linear Model. However, the potential of the ZINB model in the microbiome field has not yet been fully utilized in practice. Microbial count data are more complex than other scenarios, and our model captures this complexity. Our study is the first to discuss the effects of zero inflation and the degree of overdispersion in microbiome data on the results of model solutions. Finally, our work successfully applied it to a real multi-ethnic cohort study.

## INTRODUCTION

Humans harbor a extremely complex gut microbiota whose composition diversifies between different population. It plays a critical role in performing essential physiological functions for the life and health [1, 2]. In recent decades, the development of High-Throughput Sequencing (HTS) technology has greatly advanced the progress of microbiology, shifting the perspective of microbiologists from culture to non-culture, and opening up new ways for us to understand trillions of microorganisms [3]. Typically, microbiome data is obtained by amplifying and sequencing variable regions of the 16S rRNA gene from a few samples of interest, using HTS techniques. The results of dividing 16S rRNA genes into Operational Taxonomic Units (OTU) or Amplicon Sequence variants (ASV) were summarized into multidimensional vectors of OTU/ASV counts for all samples. These count data provide information about the microbial composition and distribution of each sample with high resolution. Due to the continued explosion of HTS data on microbial communities, statisticians intend to develop a variety of methods to accurately characterize microbiome data, especially for complex microbial ecosystems, but the current statistical and analytical methods used remain challenging.

At first, statistical methods were used to explain differences in environmental factors by comparing the level of significance between different groups (environmental factors). However, since each covariate can only be analyzed separately as a categorical variable, these methods cannot handle the correlation between covariates well. Furthermore, evaluating the contribution of each covariates by regression analysis is a great strategy, which focuses on the comparison of microbiome community, such as the comparison of multi-taxa [4, 5], core microbiomes or enterotype, as well as the ratios(e.g. Firmicutes/Bacteroidetes ratio [6, 7]) of certain bacteria. However, these methods do not characterize differentially abundant species well, resulting in difficulties in clinical trials, mechanism validation, and biological replication. Another approach take each individual bacteria as a dependent variable for different subject groups or conditions, interrogates one-by-one. Unfortunately, one deficiency of these methods is that the optimization for microbiome count datasets has been ignored.

Current literature shows that due to the over-dispersion, zero-inflation, and fluctuating library size of the microbial count data [8, 9], the analysis of microbial data is complicated. Usually, in many microbiome data analyses, Gaussian distributions are used to simplify the calculation, without considering properties of the microbiome count data. In fact, using a Gaussian distribution model for regression analysis can lead to unreliable coefficient estimates [10], thereby undermining the reliability and accuracy of the model. The existing literature shows that negative binomial regression models are more suitable for a variety of data that exhibit a larger variance or/and overdispersion [11, 12, 13, 14], whereas the Hurdle [15] and the zero inflation counting model, has been found to be effective when processing count data having an excessive number of zero. Recently, based on the advantages and disadvantages of the above research models, ZINB models were proposed by combining negative binomial model and zero inflation counting model. ZINB models are applicable when there is interest in a model for latent taxa corresponding to a impressionable microbiota at risk for the sequencing/PCR amplifying condition under study with counts generated from a negative binomial distribution and a non-impressionable microbiota that provides only zero counts. However, this methods are challenging to implement in real scenarios due to complex coding and theoretical obscurity.Indeed, it was sporadically applied to some specific research.

A growing number of studies have confirmed that some key taxa of the microbiome are closely associated with physical indicators[16, 17, 18] of the body, such as height, weight, age and BMI, which can be used as measures of human health. However, despite advancements in understanding the gut microbiota, it is unclear which aspects of ethnicity and geography [19, 20], whether culture-related activities (lifestyle) or genetics, underlie its observed association with the microbiome. How to accurately identify which taxa in the microbiome are closely related to indicators of physical development is valuable as microbial markers of local child growth [21, 22, 23]. In practice, due to the complexity of geographical factors and ethnic factors, as well as the difficulty of sampling, it is difficult to carry out the research accurately.

In our study, through high-throughput sequencing of the V1-V3 region of the 16S RNA gene, we obtained intestinal microbiome data of four healthy children cohorts (585 preschool and school-age children) from two ethnic groups in Yili Prefecture, Xinjiang, western China, who resided spatially isolated and whose parental groups rarely intermarry across ethnic groups [24]. Our research focused on understanding the normal developmental trajectories of the gut microbiota in growing children and the microbial taxa associated with body index,thereby promoting an understanding of the interactions between ethnic and geographical factors in the child’s setting. To achieve this goal, we optimized a Zero-Inflated Negative Binomial (ZINB) model aiming to addresses the complexity associated with zero inflation, overdispersion, and multivariate structure of gut microbiome data described above. It is expected that this model can be accurately applied to microbiome data of multiple child cohorts. Our model has a unique strength in that it can not only identify differentially abundant taxa among multiple subject groups but also quantify the associations between taxa and covariates, such as body index. In terms of the analysis process, this study has its advantages in the following treatment: 1) methods for analyzing microbiome data involving Bayesian variable selection strategies; 2) constructing Bayesian regression based on the generalized linear model framework, which combines zero expansion, over-dispersion and multiple correlation structures; 3) designing a analysis pipeline for microbial data with zero-expansion counting, including analysis of response variables one by one and taxonomic comparison, with special emphasis on the association between the microbiome data and covariates.

### Zero-inflated count data regression models

#### Sampling model

The HTS data were analyzed to obtain a high-dimensional counts matrix (OTU/ASV table). This represents every count of taxonomy bacterial species that the host individual contains. In this study, the count regression model is the preferred model of analysis. We assume that non-negative integer counts *Y*_*ij*_ are observed for OTU/ASV j in sample i, j *∈* {0, …, J} and i *∈* {1, …, I}, and are organized in a I × J table, Y = [*Y*_*ij*_]. In the zero-inflated negative binomial regression model, the crucial objectivates are to make sure the significant factors could influence the *Y*_*ij*_, and to determine the extent of the effect of potential host and environmental factors on the count of non-negative integer counts *Y*_*ij*_. Specifically, we assume that each *y*_*j*_ = *Y*_*ij*_ conforms to a separate zero-inflated count distribution, follows as:

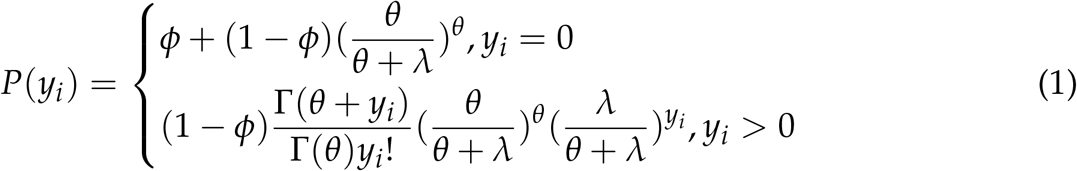

where *ϕ ∈*[0,1] is the weight of extra zeros generated from the sampling count missing, including biological zeros and technical zeros [25]. Thus, the zeros of OTU/ASV counts include both fixed zeros from absence of species, and random zeros from unknown group membership of present subjects. Although the random zeros generated from present subjects have intrinsic interest, their mixture has been treat as a extremely convenient construct that came into fitting a statistical distribution of excess zeros in bacterial counts. *λ* represents an average of *Y*_*ij*_, and the dispersion parameter *θ*^−1^ is adjusted to a negative binomial distribution (NB)(so the variance is 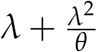) under the Poisson distribution (the variance is *λ*).

Thus,the joint distribution across all i individuals in the sample based on Eq (2) is:

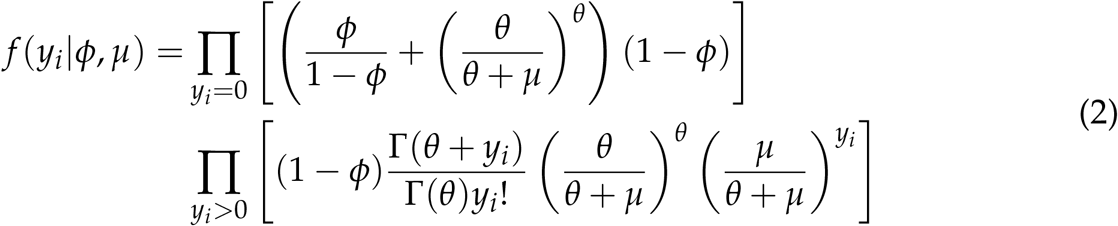

where *ϕ* = (*ϕ*_1_, *ϕ*_2_, *ϕ*_3_, …, *ϕ*_*j*_), *μ* = (*μ*_1_, *μ*_2_, *μ*_3_, …, *μ*_*j*_), and *θ* = (*θ*_1_, *θ*_2_, *θ*_3_, …, *θ*_*j*_) are dispersion parameter that is assumed not to depend on covariates.

In the generalized linear models, *log*(*μ*) and *logit*(*ϕ*) are transformations that successfully linearize poisson means and Bernoulli probabilities through logit model [26] as follows Equation (3):

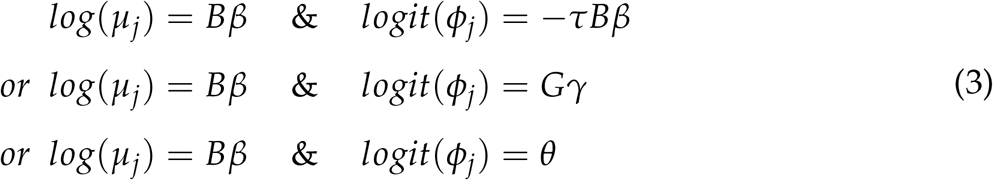

where B are the designed covariates matrices and the *β* are the regression coefficient vectors. *τ* is defined as real-valued shape parameter [27], which implies that *ϕ* = (1 + *λ*^*τ*^)−^1^.

### Likelihood function establishment

In this situation, the overall mean of specific bacteria *μ* = *E*[*y*_*i*_] is the primary interest.

The vector parameter of *log*(*μ*)’s with the intercept *β*_0_ included denoted by *β* = (*β*_0_, *β*_1_, …, *β_j_*−1)^*′*^ represents the same overall effect of covariates on species counts increment as in ZINB regression. In other words, exp(*β*) represents the multiplicative increase in *log*(*μ*) count for species in the overall counts corresponding to a one-unit increase in the covariates matrices B.

*β* in a maximum likelihood framework as follows:

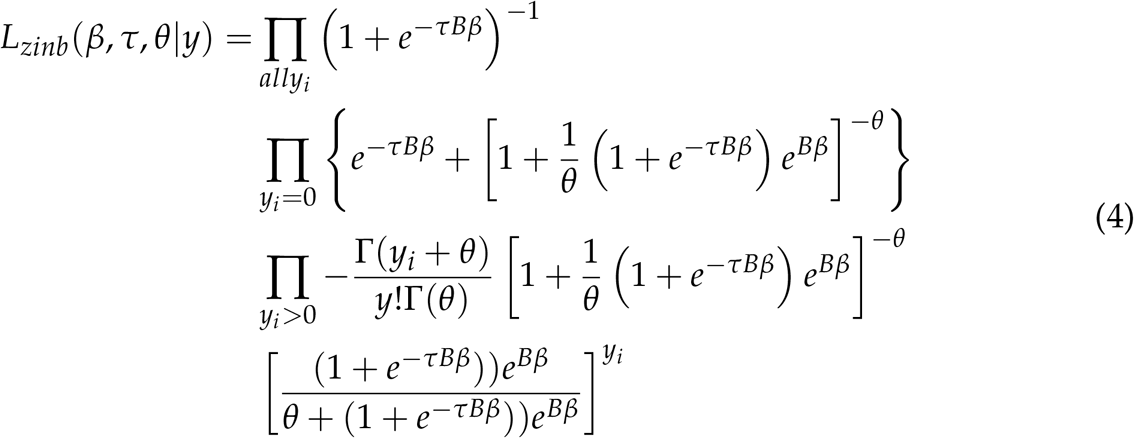

The log-likelihood of the ZINB model is:

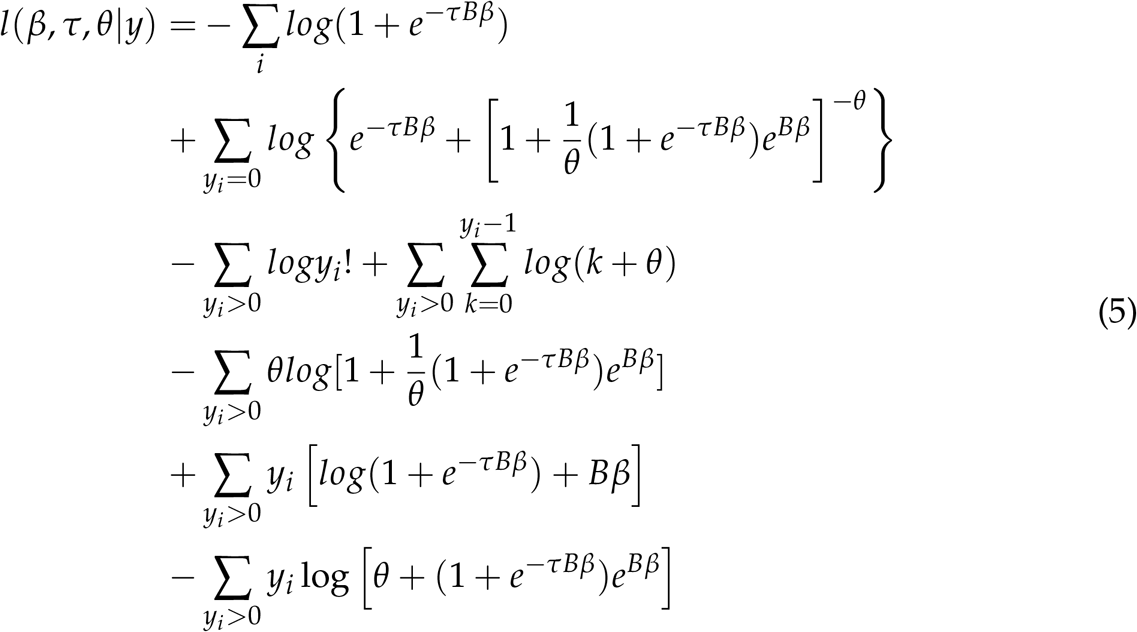

### Estimation of the distinct-parameters ZINB regression model

We need to specify a prior distribution for parameters in the model to obtain a Bayesian estimation of the unknown parameters in the ZINB model. A great result depends on establishing a good initial prior information as well as a formulation of an informative prior distribution [28]. In this model, we assumed that the overdispersion parameter *θ* and shape parameter *τ* follow the Gamma distribution(*θ* ∼ dgamma (a = 0.001, b = 0.001)) and model coefficient parameter *β* follows the Normal distribution(0, 10^−6^). Thus, the prior distribution formulation is as the following:

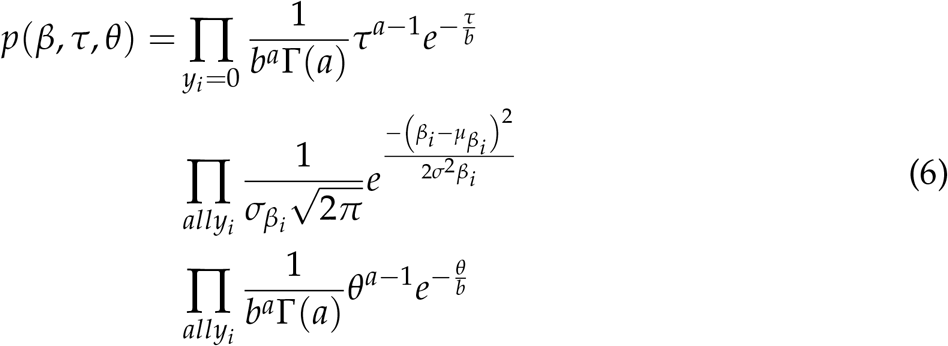

The posterior distribution of parameters can be obtained by combining the likelihood function (Eq (5)) with the prior function (Eq (6)), as:

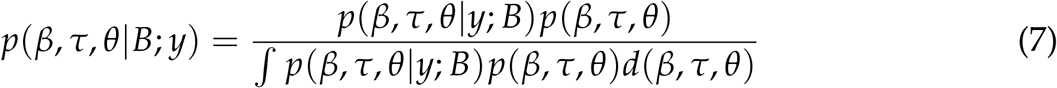

We used standard Markov Chain Monte Carlo (MCMC) methods, which are available in JAGS using runjags package [29] in R software. The estimated effect size will be determined by the Bayesian estimated average. We will assess the uncertainty of the estimated effect size with credible intervals.

## Methodologies

### Design and Study Population

This study was conducted in accordance with the guidelines of the Declaration of Helsinki [30] and was approved by the Ethics Committee of the First Affiliated Hospital of Shihezi University Medical School (2017-117-01). For school-aged children, we communicated with local authorities and school principals to get permission to sample. In addition, the researchers trained the teachers, and the teachers publicized the purpose and significance of our research. At the same time, informed consent forms were issued, which the students take home for parents to decide whether to sign or not. The following days, the participants had their height and weight measured and were inquired about their date of birth and recent dietary habits. After the survey, the participants were supplied with a stool sampler, ice bag, and aseptic bag, and were given comprehensive guidance on how to collect and preserve the samples. Participants are required to send their stool samples back to school as soon as possible, ideally within 1 to 2 days. The criteria for inclusion in the study were as follows: (i) school-aged children, (ii) ethnic minorities, and both parents are ethnic minorities, (iii) able to provide informed consent, (iv) no antibiotics or other medications that could affect the composition of gut microbes have been used in the past 6 months, (v) be willing to provide regular stool samples as required by the study. The criteria for exclusion in the study were as follows: (i) individuals with severe digestive disorders (e.g. Crohn’s disease, ulcerative colitis) or other chronic diseases (e.g., diabetes, heart disease), (ii) recent (within 6 months) use of antibiotics or other drugs that affect the gut microbiota, (iii) iIndividuals who are unable to provide informed consent.

We obtained a total of 776 samples from June 8, 2021, of which only 585 were used in this study. The survey was a population-based cross-sectional study that included four different geographic locations(4 primary schools, 2 kindergartens), two distinct ethnic groups (Uyghur and Kazakh), and an age range between 4 and 15 years. All the recruits were from rural locations. Despite the presence of numerous ethnic groups in Xinjiang, only the samples from the Uyghur and Kazakh populations we gathered fulfilled our study criteria (The sample size of other ethnics are too small …).

### Stool sample collection and transportation

The participants had their height and weight measured and were inquired about their date of birth and recent dietary habits. After the survey, the participants were supplied with a stool sampler, ice bag, and aseptic bag, and were given comprehensive guidance on how to collect and preserve the samples. The instructions stated that the feces are placed in stool sampler should be placed in a sterile sampling bag with an ice pack in the participant’s home refrigerator (-20 to -18 ^*°*^C) until sample collection. Participants are required to send their stool samples back to school as soon as possible, ideally within 1 to 2 days. Collected samples were stored in a car refrigerator (-20 to -18 ^*°*^C) and then were transported to the research laboratory (Shihezi University food Biotechnology Research Center). The samples were transported to the research laboratory and then stored in freezers at -80 ^*°*^C until they were ready for further processing.

### DNA extraction and 16S rRNA Sequencing

Total genomic DNA was extracted from faeces samples using the E.Z.N.A.® Soil DNA Kit (Omega Bio-tek, Norcross, GA, U.S.) according to manufacturer’s instructions. Concentration and purity of extracted DNA was determined with TBS-380 and NanoDrop2000, respectively. DNA extract quality was checked on 1% agarose gel. DNA extract was fragmented to an average size of about 400 bp using Covaris M220 (Gene Company Limited, China) for paired-end library construction. Aired-end library was constructed using NEXTFLEX Rapid DNA-Seq (Bioo Scientific, Austin, TX, USA). Adapters containing the full complement of sequencing primer hybridization sites were ligated to the blunt-end of fragments. Paired-end sequencing was performed on Illumina Novaseq 6000 (Illumina Inc., San Diego, CA, USA) at Majorbio Bio-Pharm Technology Co., Ltd. (Shanghai, China) using NovaSeq Reagent Kits according to the manufacturer’s instructions (www.illumina.com). The extracted DNA was used as a template for PCR amplification of the V1-V3 region of bacterial 16S rRNA genes by HiSeq sequencing. In brief, samples amplified with “27f-YM” used forward primer YM-27F (5’-AGRGTTYGATYMTGGCTCAG-3’) [31] and a reverse primer 534R (5’-ATTACCGCGGCTGCTGG-3’). The PCR conditions were as follows: start at 95 ^*°*^C for 4 min, 30 cycles denaturing at 95 ^*°*^C for 30 s, annealing at 72 ^*°*^C for 50 s, and final extension at 72 ^*°*^C for 10 min.

### Bioinformatics and biostatistics

Paired-end sequencing reads were processed using a usearch (v11.0.667_i86linux64) pipeline [32]. The entire workflow includes merging sequences, trimming barcodes and primers, conducting sequential quality control, and generating zero-radius operational taxonomic units (zOTUs), aka amplicon sequence variants (ASVs). Taxonomy of the ASVs were then predicted using the SINTAX algorithm [33], with the settings strand both and “sintax_cutoff” 0.8 using the SILVA SSU rRNA 138 database [34]. Finally, an abundance table was generated using the “otutab” command by mapping the zOTUs obtained from filtered read.

We will use rarefaction to standardize the sequencing depth across samples. The rarefaction depth will be set to 5000 sequences for each sample. For analyzing the microbiota alpha and beta diversity, Shannon index, Bray-Curtis distance were estimated using vegan R package [35].

### Weight, Height and BMI for age Z-scores

We separately calculated weight, height and BMI (kg/*m*^2^) z-scores (z-BMI) standardized on age and sex as indicated by WHO Growth Reference (2006) charts using the zscorer (v.0.3.1) [36] package in R. The z-BMI standard deviation is usually used for descriptive statistics to classify individuals as follows: Normal (>-2 sd and ≤ 2 sd), overweight (>2 sd and ≤ 4 sd), and obese (>4 sd).

### STORMS Checklist

This study has been completed according to the STORMS Checklist (DOI: https://doi.org/10.5281/zenodo.11127032).

## Results

### Simulation Data generation and description

In the simulation studies, it is important to have a clear understanding of the properties of both covariates and response variables before conducting the simulations. It is assumed that the microbial population follows a zero-inflated negative binomial distribution, and it has been established that there are six covariates that may have an impact on it. Some of these covariates may have interactions, and therefore we analyze them separately.

#### Response variable

- **count** – bacterial counts for ASV table follows a negative binomial distribution.

#### Covariates

- **Age** – a discrete variable with a range of (3.0 ∼ 14.0)
- **Sex** – a classified variable with two sex male an female.
- **Ethnic** – classified variable with two enthic Uighur (U) or Kazakh (K)
- **Geography** – classified variable with two location H or W.
- **Height** – a discrete variable with a range of (0.90 ∼ 1.50)
- **Weight** – a discrete variable with a range of (17.0 ∼ 27.0)

#### Eliminate effects of interaction

- **BMI** - a discrete variable equal to Weight/(Height)^2^. To eliminate the interaction both Weight and Height.
- **BMIAZ** - BMI for Age Z-scores, to eliminate the interaction of Weight, Height and Age.
- **HAZ** - Height for Age Z-scores,
- **WHZ** - Weight for Height Z-scores,
- **WAZ** - Weight for Age Z-scores,

The item “Eliminate effects of interaction” in the above list refers to eliminating the interactions by differentiating the Z-scores of the body index. The properties of covariates, such as their distribution and range of settings in which they are measured, are explained in detail.

### Z-score processing is better to eliminate interaction between covariables

In the experiment, the interactions brought about by covariates are often uncertain. We try to eliminate the interaction between covariates by using several processing method as the Table 1 summarizes. We interact with generalized linear models (GLM) for covariables that we suspect may interact, such as ethnic and geography (maxGVIF = 125.87); height and weight (maxGVIF = 125.76). The experiment results show that GLM don’t eliminate interactions very well. Nextly, we tried to replace weight and height with BMI, GVIF value decreased significantly (maxGVIF = 16.40). The results show that interaction is reduced significantly, but it still doesn’t reach the conditions for naive bayesian (GVIF *<* 3) [37, 38]. This can indicate that BMI index tend to change with age. Finally, we use BMI for age Z-scores index to replace height *\** weight interaction. Interaction is eliminated successfully (maxGVIF = 2.12).

**TABLE 1.**
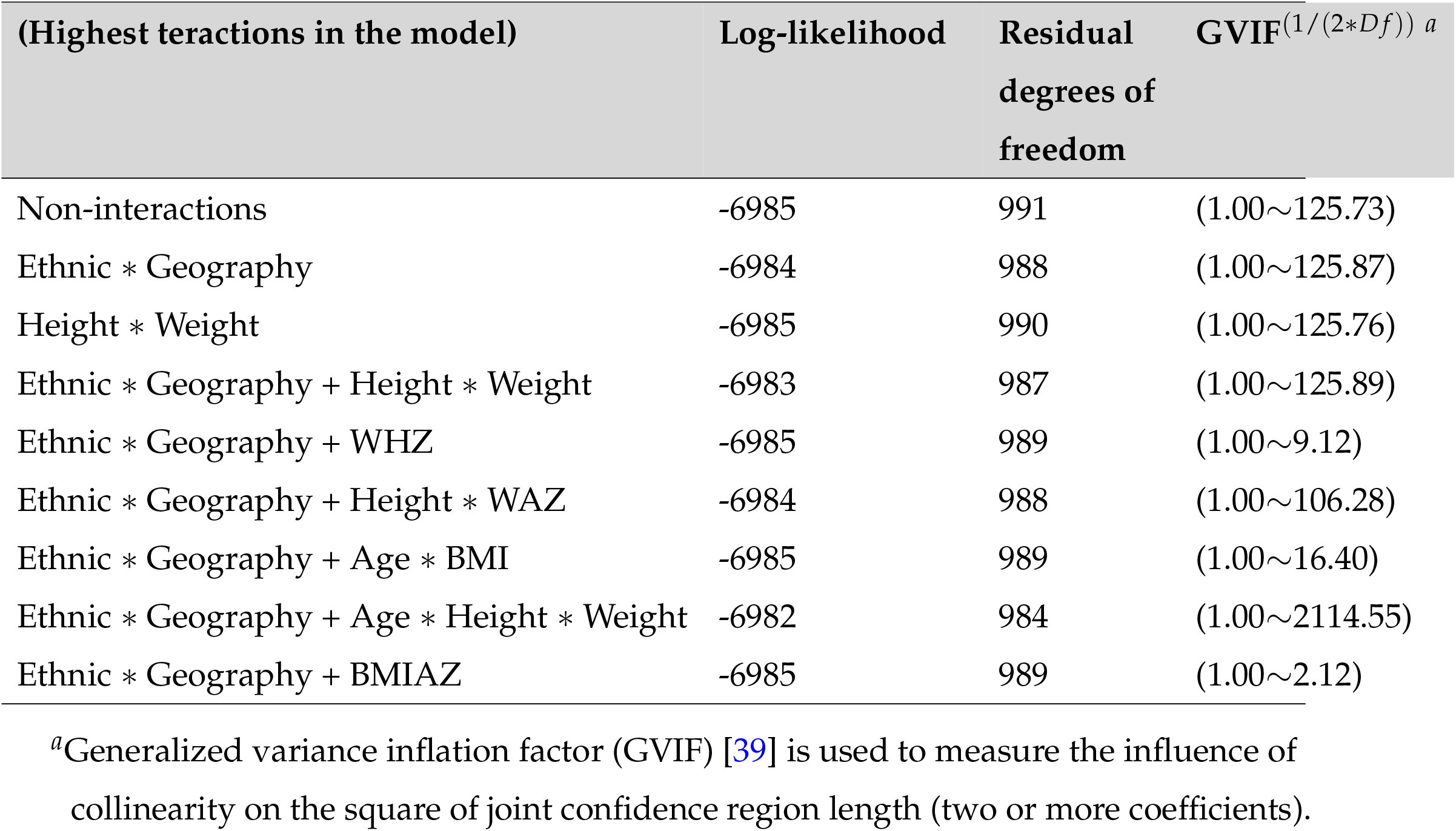
Fits of Some ZINB Models for the Number of Bacterial counts.

### The simulation experiments

To examine the finite sample properties of ZINB model, simulations were performed the count data regression. Let *Y*_*i*_ be the bacterial counts for the *i*th Sample, *x*_*i*1_ is age (mean = 8, sd = 2.5); *x*_*i*2_ ethnic (binary), and *x*_*i*3_ is the BMIAZ (mean = 0, sd = 1). Data were generated from the R, and all the variates follow the ZINB model’s distribution. For the parameter, i assume *β* = (0, -0.5, 1.0, 1.5), *γ* = (0, -0.25, 0.5, 0.75) and *θ* = 1. Thus, the *τ* = 0.5 and the zero-propotion = 0.25.

Each simulation scenario consisted of the following steps:

1. Take random samples from each dataset with sizes of 100, 200, 500 and 1000;
2. Set iteration parameters as adapt = 10000, burning = 2000, sample = 2000;
3. Fit four models:
  (a) Hurdle models [40]
  (b) ZINB(*τ*) models
  (c) MZINB models
  (d) ZINB(inter) models
4. Record all simulation parameters and estimate their deviations.

Table 2 reports the estimated parameter values and standard errors. Where the estimation of *μ* in the five models are same, but difference in estimation of *ϕ*. Hurdle and MZINB model followed the link function *logit*(*ϕ*) = −*γ*_0_ + *γ*_1_ *\* x*_1_ + *γ*_2_ *\* x*_2_ + *γ*_3_ *\* x*_3_, ZINB(*τ*) model followed *logit*(*ϕ*) = −*τ*(*β*_0_ + *β*_1_ *\* x*_1_ + *β*_2_ *\* x*_2_ + *β*_3_ *\* x*_3_), and the Equation *logit*(*ϕ*) = *θ* guides ZINB-inter model with *θ* belong to gaussian distribution. In general, ZINB(*τ*) gives the least biased estimates with covariate effection, but intercept is not a good estimate. For simulations with a sample size of less than 500, all models are not very good estimates, and bias decreases with increasing sample size. All bayesian method are superior to maximum likelihood method (Hurdle) for solving complex models.

**TABLE 2.** Fits of Some ZINB Models for the Number of Bacterial counts.

| Parameter | n | Hurdle | MZINB | ZINB-inter | ZINB($\tau$) |
| --- | --- | --- | --- | --- | --- |
| $\beta_0$ | 100 | -0.11(0.23) | -0.23(0.19) | -0.26(0.18) | 0.24(0.16) |
|  | 200 | -0.23(0.17) | -0.32(0.11) | -0.31(0.14) | -0.24(0.11) |
|  | 500 | 0.06(0.11) | -0.04(0.11) | -0.17(0.14) | 0.17(0.07) |
|  | 1000 | -0.01(0.08) | -0.03(0.07) | -0.12(0.11) | 0.14(0.06) |
| $\beta_1$ | 100 | -0.50(0.06) | -0.51(0.06) | -0.52(0.06) | -0.47(0.04) |
|  | 200 | -0.48(0.04) | -0.49(0.03) | -0.50(0.04) | -0.43(0.03) |
|  | 500 | -0.52(0.03) | -0.53(0.02) | -0.53(0.02) | -0.47(0.02) |
|  | 1000 | -0.47(0.02) | -0.47(0.02) | -0.47(0.02) | -0.45(0.01) |
| $\beta_2$ | 100 | 1.25(0.25) | 1.13(0.23) | 1.17(0.23) | 1.30(0.21) |
|  | 200 | 1.12(0.17) | 1.04(0.12) | 1.04(0.16) | 1.11(0.05) |
|  | 500 | 0.84(0.11) | 0.85(0.10) | 0.93(0.11) | 0.99(0.07) |
|  | 1000 | 1.05(0.08) | 0.96(0.09) | 1.05(0.08) | 1.05(0.03) |
| $\beta_3$ | 100 | 1.52(0.12) | 1.53(0.10) | 1.56(0.11) | 1.42(0.10) |
|  | 200 | 1.63(0.09) | 1.61(0.08) | 1.59(0.07) | 1.48(0.07) |
|  | 500 | 1.48(0.06) | 1.54(0.06) | 1.55(0.06) | 1.46(0.04) |
|  | 1000 | 1.47(0.05) | 1.49(0.04) | 1.46(0.04) | 1.48(0.03) |
| $\tau$ | 100 | | | | 0.51(0.15) |
|  | 200 |  |  |  | 0.45(0.08) |
|  | 500 |  |  |  | 0.53(0.06) |
|  | 1000 |  |  |  | 0.47(0.04) |
| $\theta$ | 100 | 0.97 | 0.55(0.09) | 0.54(0.08) | 1.22(0.27) |
|  | 200 | 0.86 | 0.51(0.05) | 0.44(0.04) | 1.14(0.15) |
|  | 500 | 1.06 | 0.98(0.13) | 0.78(0.22) | 1.08(0.12) |
|  | 1000 | 0.98 | 0.96(0.09) | 0.75(0.23) | 1.05(0.08) |
| $\gamma_0$ | 100 | -0.49(0.23) | -84(93) | -806(614) | |
|  | 200 | -0.49(0.15) | -74(55) | -812(610) |  |
|  | 500 | -0.60(0.11) | -1.59(0.61) | -400(585) |  |
|  | 1000 | -0.55(0.07) | -1.55(0.32) | -398(573) |  |
| $\gamma_1$ | 100 | -0.38(0.08) | -10.1(8.7) | | |
|  | 200 | -0.35(0.05) | -7.3(7.5) |  |  |
|  | 500 | -0.36(0.04) | -0.03(0.12) |  |  |
|  | 1000 | -0.33(0.02) | -0.06(0.05) |  |  |
| $\gamma_2$ | 100 | 0.75(0.34) | -105.4(97.1) | | |
|  | 200 | 0.51(0.21) | -63.2(61.7) |  |  |
|  | 500 | 0.79(0.15) | -0.55(0.36) |  |  |
|  | 1000 | 0.75(0.10) | -0.09(0.23) |  |  |
| $\gamma_3$ | 100 | 1.04(0.19) | -4.52(8.42) | | |
|  | 200 | 0.98(0.11) | -3.70(9.33) |  |  |
|  | 500 | 1.10(0.08) | -0.02(0.28) |  |  |
|  | 1000 | 0.92(0.06) | 0.14(0.15) |  |  |
| zero-propotion | 100 | 0.62(0.25) | 0(0) | 0.01(0.00) | 0.21(0.04) |
|  | 200 | 0.55(0.26) | 0.01(0.00) | 0.01(0.00) | 0.24(0.03) |
|  | 500 | 0.61(0.26) | 0.15(0.04) | 0.08(0.03) | 0.21(0.02) |
|  | 1000 | 0.60(0.25) | 0.17(0.03) | 0.09(0.04) | 0.25(0.01) |

The table shows the simulation results of different parameters of the four models (Hurdle, MZINB, ZINB-inter and ZINB( $\tau$ )), where n represents the number of Bayesian samples. The blank part indicates that the model does not contain this parameter.

**TABLE 3.** The average and standard deviation among four models in the simulation study.

| Overdispersion<br>($1/\theta$) | n | Hurdle<br>$1/\theta$ | ARB | MZINB<br>$1/\theta$ | ARB | ZINB-inter<br>$1/\theta$ | ARB | ZINB($\tau$)<br>$1/\theta$ | ARB |
| --- | --- | --- | --- | --- | --- | --- | --- | --- | --- |
| 0.5 | 100 | 0.73(0.23) | 4.38(0.69) | 0.66(0.18) | 2.59(1.69) | 0.56(0.11) | 3.35(0.55) | 0.48(0.20) | 3.35(0.55) |
|  | 200 | 0.41(0.17) | 2.57(0.58) | 0.86(0.17) | 2.75(1.92) | 0.67(0.21) | 3.32(0.37) | 0.37(0.20) | 3.12(0.55) |
|  | 500 | 0.47(0.11) | 2.71(0.11) | 0.47(0.14) | 3.29(1.55) | 0.47(0.14) | 3.40(0.37) | 0.39(0.19) | 3.45(0.58) |
|  | 1000 | 0.37(0.08) | 2.86(0.07) | 0.39(0.12) | 3.70(1.52) | 0.47(0.11) | 3.02(0.36) | 0.31(0.20) | 3.74(0.58) |
| 1 | 100 | 1.78(0.23) | 0.62(0.17) | 3.46(2.70) | 3.46(2.70) | 0.58(0.34) | 3.30(0.21) | 1.68(0.85) | 3.63(2.74) |
|  | 200 | 1.18(0.17) | 0.61(0.07) | 3.21(1.56) | 3.64(0.11) | 0.81(0.14) | 3.64(0.11) | 1.43(0.41) | 3.25(1.55) |
|  | 500 | 1.05(0.11) | 0.63(0.14) | 3.74(0.77) | 3.56(0.07) | 0.98(0.13) | 3.56(0.07) | 1.22(0.20) | 3.77(0.76) |
|  | 1000 | 1.00(0.08) | 0.74(0.07) | 3.29(0.78) | 3.59(0.06) | 0.94(0.11) | 3.59(0.06) | 1.06(0.20) | 3.32(0.78) |
| 2 | 100 | 1.96(0.23) | 2.68(0.69) | 4.13(1.66) | 1.35(0.90) | 0.92(0.74) | 2.04(0.31) | 1.50(0.64) | 1.38(0.93) |
|  | 200 | 1.87(0.17) | 2.40(0.11) | 6.05(2.71) | 1.69(0.65) | 1.02(0.54) | 2.45(0.11) | 1.59(0.17) | 1.65(0.76) |
|  | 500 | 1.30(0.11) | 3.33(0.11) | 3.79(1.00) | 2.76(0.58) | 1.39(0.14) | 3.21(0.07) | 1.62(0.31) | 2.73(0.60) |
|  | 1000 | 1.72(0.08) | 3.59(0.07) | 3.67(0.64) | 2.69(0.40) | 1.69(0.11) | 3.15(0.06) | 1.81(0.19) | 2.72(0.40) |
| 5 | 100 | NA | NA | NA | NA | NA | NA | NA | NA |
|  | 200 | NA | NA | NA | NA | NA | NA | NA | NA |
|  | 500 | 2.27(0.11) | 3.50(0.11) | 19.12(4.20) | 7.64(2.75) | 1.55(1.20) | 2.89(0.07) | 7.91(4.42) | 1.53(0.54) |
|  | 1000 | 2.94(0.08) | 2.95(0.34) | 16.12(4.52) | 4.58(1.99) | 2.51(1.11) | 2.74(0.06) | 7.16(4.21) | 1.68(0.45) |

The $1/\theta$ in the first column is the parameter set by the model, and the $1/\theta$ in the following columns is the simulation result. ARB represents the average difference between the actual value of each fitting result and the real value. "NA" indicates that the mode can not calculate the parameter result (Slicer stuck at value with infinite density).

Further more, we evaluate the effects of overdispersion generation on the model. We follow the parameter Settings mentioned above and estimate the model robustness by changing the overdispersion parameters. We set four overdispersion parameters (1/*θ* = 0.5, 1, 2, 5) to simulate the model’s response to excessive dispersion. Table 4 shows the four models estimated accuracy by reporting the Average Relative Bias (SD) and overdispersion estimation. The results indicate that both the Hurdle and the ZINB-inter models underestimated overdispersion, while the MZINB model overestimated overdispersion vastly. The ZINB(*τ*) model provides more accurate estimates than other models in cases of over-dispersion, as indicated by its lower ARB values. Furthermore, in instances of extreme overdispersion (with 1/*θ* = 5), all the models will have unestimable parameters (Slicer stuck at value with infinite density) for sample sizes below 500. Aiming at these problems, Zitzmann et al. [41] reported and raised a question as to whether bias is the most important criterion to consider when the sample size is small. We hold the opinion that large-sample is the most effective way.

### Real Data Application

In our study, information of gut microbiome and body development index were obtained from a total of 585 healthy preschool and school-age children from Ili district of the westernmost part of Xinjiang, China. They were recruited from four different villages (4 primary schools and 3 kindergartens) and mainly belong to two ethnic groups, Uyghur and Kazakh, as shown in Fig. 1a. Overall, the cohort of children in each village was overwhelmingly dominated by one of the ethnic groups, every cohort have an even ratio of girls to boys (Table S1 at https://doi.org/10.5281/zenodo.11127871). The gut microbiota of the children was examined through 16s amplicon sequencing. After removing noise from the 585 samples, we obtained 10,065 ASVs (Table S2, S3 at https://doi.org/10.5281/zenodo.11127871). Firstly, we figured up measures of gut bacterial alpha diversity, including Shannon’s index. We did not observe marked ethnic differences in Shannon’s index (P *>* 0.05, wilcox test), only measure from H and S villages show slight differences in Shannon’s index (P = 0.029, Shannon’s index) as shown in the Fig. 1b. In Fig. 1c, obesity-related differences are shown (P *>* 0.05, wilcox test). Then, we performed statistics on bmiAgeZ for children cohorts of different ethnics in four villages, and the resulting date indicated the H villages children bmiAgeZ was significantly higher than other groups (P = 1.1 *×* 10^−12^, wilcox test), as shown in the Fig. 1d. Shannon also showed no difference (P *>* 0.05, Tukey’s test) in age gradient (the Fig. 1e). However, Principal Coordinates Analysis (PCoA) analysis clearly demonstrated different population community distinctions between BMI or age groups (Fig. 1f,g). According to the results of the PERMANOVA analysis, age and BMI index were found to significantly impact the demarcation of the microbial community among children cohorts (age: *R*^2^ = 0.013, P *<* 0.001; BMI: *R*^2^ = 0.006, P *<* 0.001).

**FIG 1.**
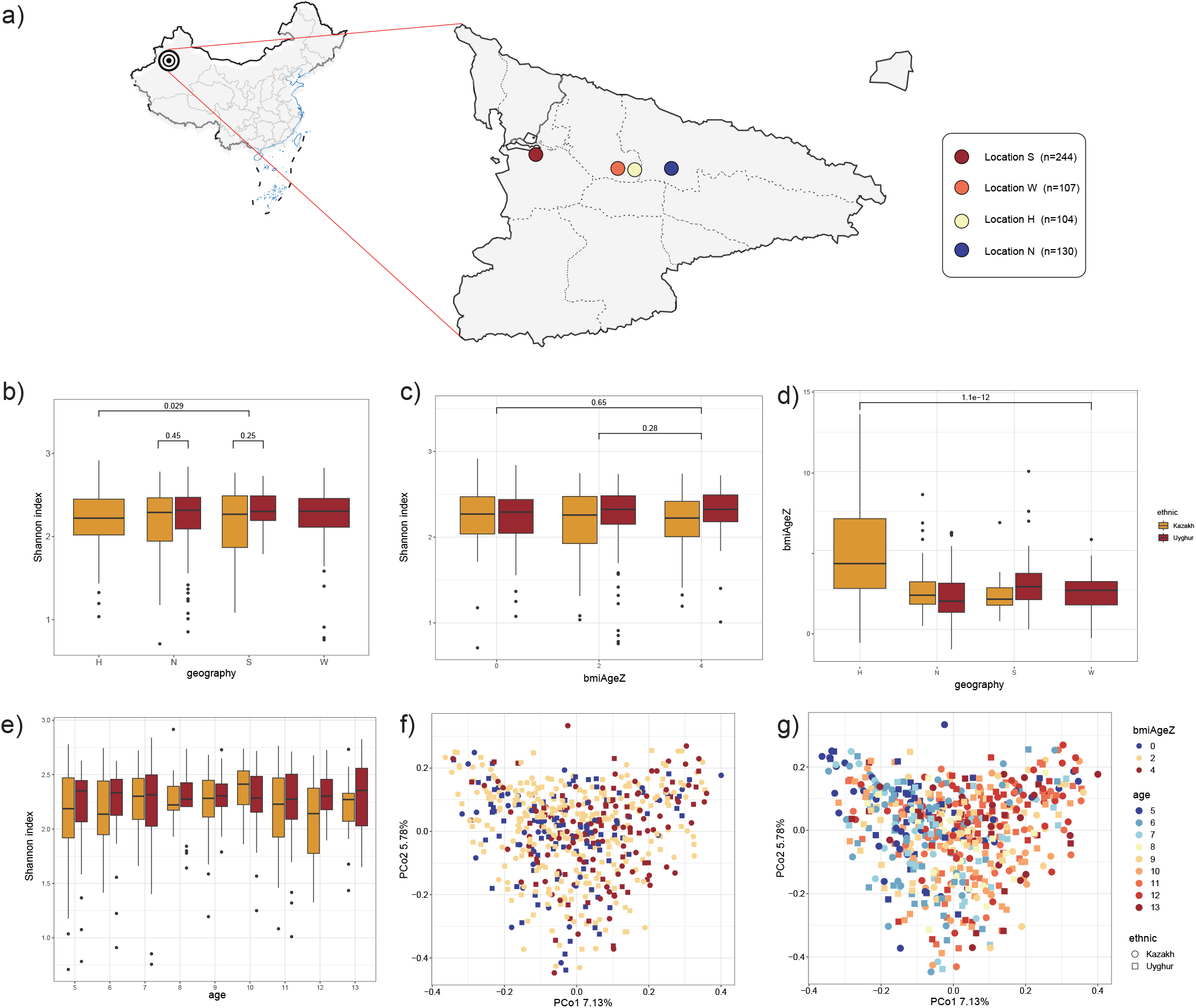
Multi-ethnic cohort of children community *α*-diversity and *β*-diversity. a) Four locations of sampling Ili Kazak Autonomous Prefecture in Xinjiang, China; b,c,e) *α*-Diversity (Shannon diversity) of microbial communities in geography, BMI and age based on V1 and V3 regions of 16S rRNA genes, respectively. The statistical method used is the Wilcoxon test performed in R and directly label the p-value above the box; d) BMI differences in the ethnic and geographical distribution; f,g) *β*-diversity across BMI and age, Principal Coordinates Analysis (PCoA) ordination of bacterial communities based on the Bray-Curtis dissimilarity.

### Interpretation of bacterial count data

In this subsection, we discuss in detail about bacterial count properties. Initially, we excluded ASVs with counts less than 15 reads due to interference for the accuracy of the calculation.

Considering a hypothesize that the zero-inflated probability (*ϕ*) and overdispersion parameters 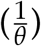 follow a Gaussian distribution, we use it to calculate their means. The formula to estimate the parameters referred to Appendix S1. We figure up the original discrete parameters of each ASV and the overdispersion parameters of the part of the negative binomial distribution. By calculating the raw data directly, we observed that only 32% of the ASV data had zero-inflated. In ASVs with zero inflation, most of the zero expansion probabilities (*ϕ*) are very high (mean = 90.2%). Furthermore, by calculating the overdispersion parameter of the negative binomial part (excluding the part of “extra-zero”), we find that the overdispersion parameter of the negative binomial part is slightly lower than that of the whole part (Fig. 2a), thus showing that zero inflation has a slight influence on the degree of data dispersion. As was shown in Fig. 2b illustrating the mean and variance for each ASV, the red solid line represents the Poisson distribution. All ASVs deviate from Poisson distribution, instead exhibit dispersion phenomena, and approximately 22.56% of ASVs showed overdispersion (with 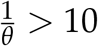) (see the Fig. 2b). This indicates that it is not advisable to choose the Poisson model for regression of microbial counts data. In Fig. 2c,d, the overdispersion and zero-inflation parameters counted at different taxonomic levels are visualized separately. We found that overdispersion parameter was the lowest at the ASV level. Moreover, the probability of zero-inflated decreasing gradually with an increase of the taxonomic rank.

**FIG 2.**
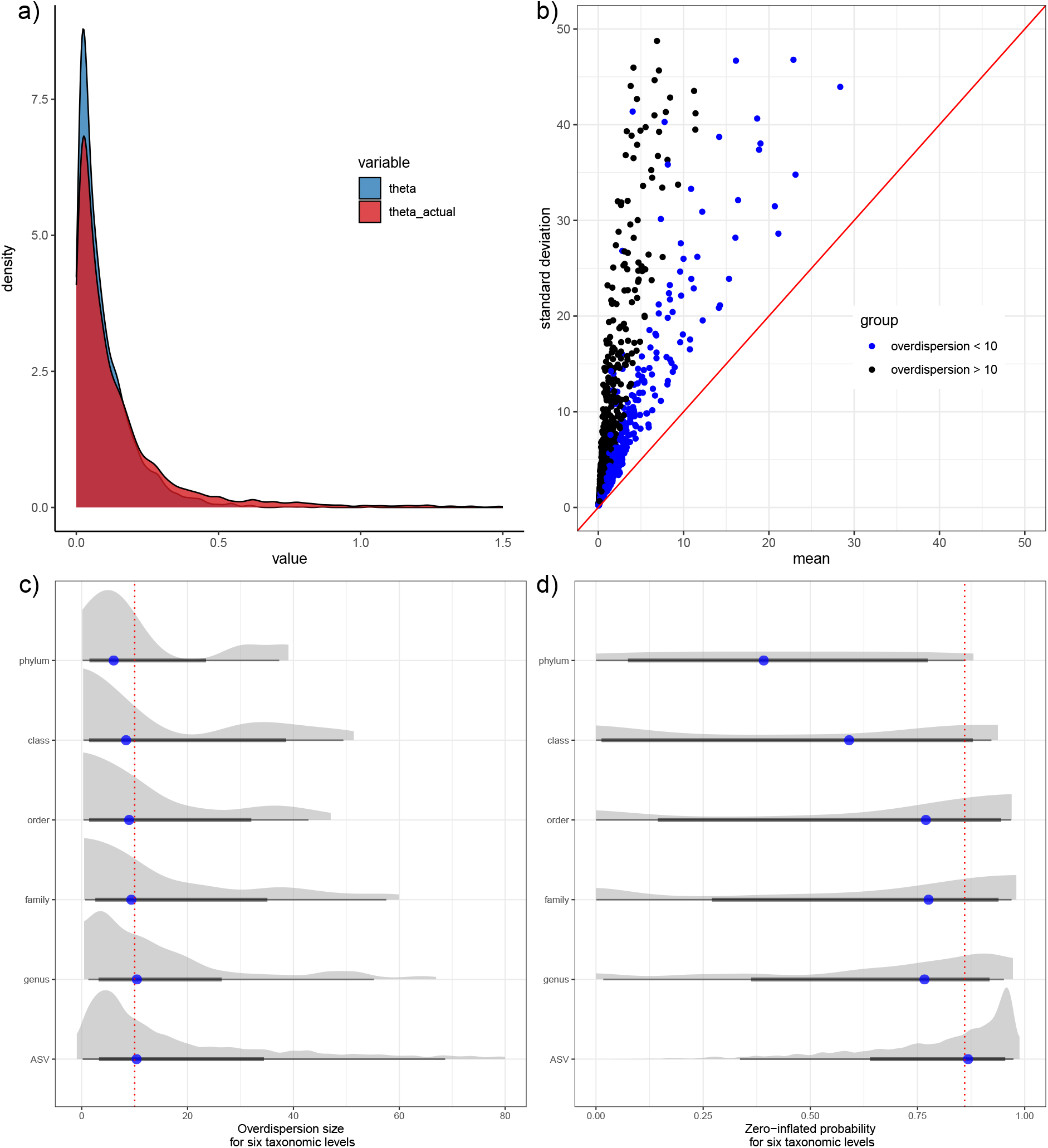
Complexity of microbiome count data. a) The discrete value of theta is estimated using the conversion equation. The blue part is the direct value and the red part is the actual value; b) Distribution of standard deviation and mean of microbial count data. Black dots indicate a value greater than 10, and blue indicates a value less than 10; c,d) Overdispersion and zero-inflation parameters of the six taxonomic levels are counted separately.

### Application to multi-ethnic children cohort study

The Bayesian ZINB model is capable of conducting integrative analysis on microbiome count data. In this subsection, we applied this model to analyze microbiome data from a multi-ethnic children cohort study. The full dataset includes 585 samples. Here, we aim to identify unique taxonomic signatures of the microbiome in children based on their age and BMI. At the species taxonomic level, the observed microbial abundance matrix Y was profiled from the gut microbiome. Before the downstream analysis, we filtered out ASVs with extremely low abundance (less than 30 observed counts in the cohort), following the suggestion in Wadsworth [42]. We obtained 2418 taxa for further analysis. To aid convergence for covariate information, center the age variable and use the BMI Z-score variable. Using a Bayesian ZINB regression model, we interrogated one-by-one each of these 2418 taxa (ASVs) and considered only the effects of sex, ethnicity, and geography, while recognizing the interaction between ethnicity and geography.

Our model seriatimly identified the age-discriminatory or BMI-discriminatory differentially abundant taxa (ASVs), and revealed the correlation between microbial groups and host physical indexes. Fig. 3a shows the posterior mean of *μY*_*ij*_ for all discriminating taxa identified. We have shown that 160 ASVs abundance remain age-discriminatory (as indicated by red dots), and 162 ASVs abundance remain BMI-discriminatory (as indicated by blue dots). For children, the average abundance of *Bacteroides, Roseburia, Faecalibacterium, Streptococcus*, and *Actinomyces* decreases as age increases [43, 44, 45]. On the other hand, *Catenibacterium, Bifidobacterium*, and *Ligilactobacillus* show an increase in average abundance with age [46]. Additionally, the average abundance of Blautia decreases with an increase in BMI for children [47], while *Actinomyces, Roseburia, Blautia, Faecalibacterium, Coprococcus, Bacteroides, Streptococcus*, and *Ligilactobacillus* show an increase in average abundance with an increase in BMI for children [48, 46, 49, 50]. Notably, *Ligilactobacillus* (ASV_4) exhibited significant impact and low variability for both age and BMI,and possesses lower levels of dispersion(1/*θ* = 2.09) and higher relative abundance (0.0372) (Fig. 3b). Some taxa are linked to both age and BMI, yet display completely opposite effect sizes, such as *Faecalibacterium* (ASV_2580, ASV_3278), *Streptococcus* (ASV_3022), *Roseburia* (ASV_3756), *Actinomyces* (ASV_ 4128) and *Bacteroides* (ASV_8238) (See the Table S4 at https://doi.org/10.5281/zenodo.11127871). In our microbita counts data, only a limited number of ASVs demonstrate zero-infated and low dispersion (Fig. 3b). Several studies have indicated a significant correlation between elevated levels of *Roseburia, Streptococcus*, and *Prevotella* with obesity. However,in other studies, conflicting outcomes were observed regarding the prevalence of *Anaerotruncus, Bacteroides, Blautia, Clostridium, Coproccocus, Faecalibacterium*, and *Ruminoccocus* in obese individuals. Indeed,the current studies show no clear conclusions about the relative abundance of these bacteria in obese or overweight people at the genus level [51, 52, 53, 54]. These results highlight the intricate nature of the gut microbial ecosystem, which means that the relative abundance of these so-called specific genera in obese individuals may vary geographically and ethnically.

**FIG 3.**
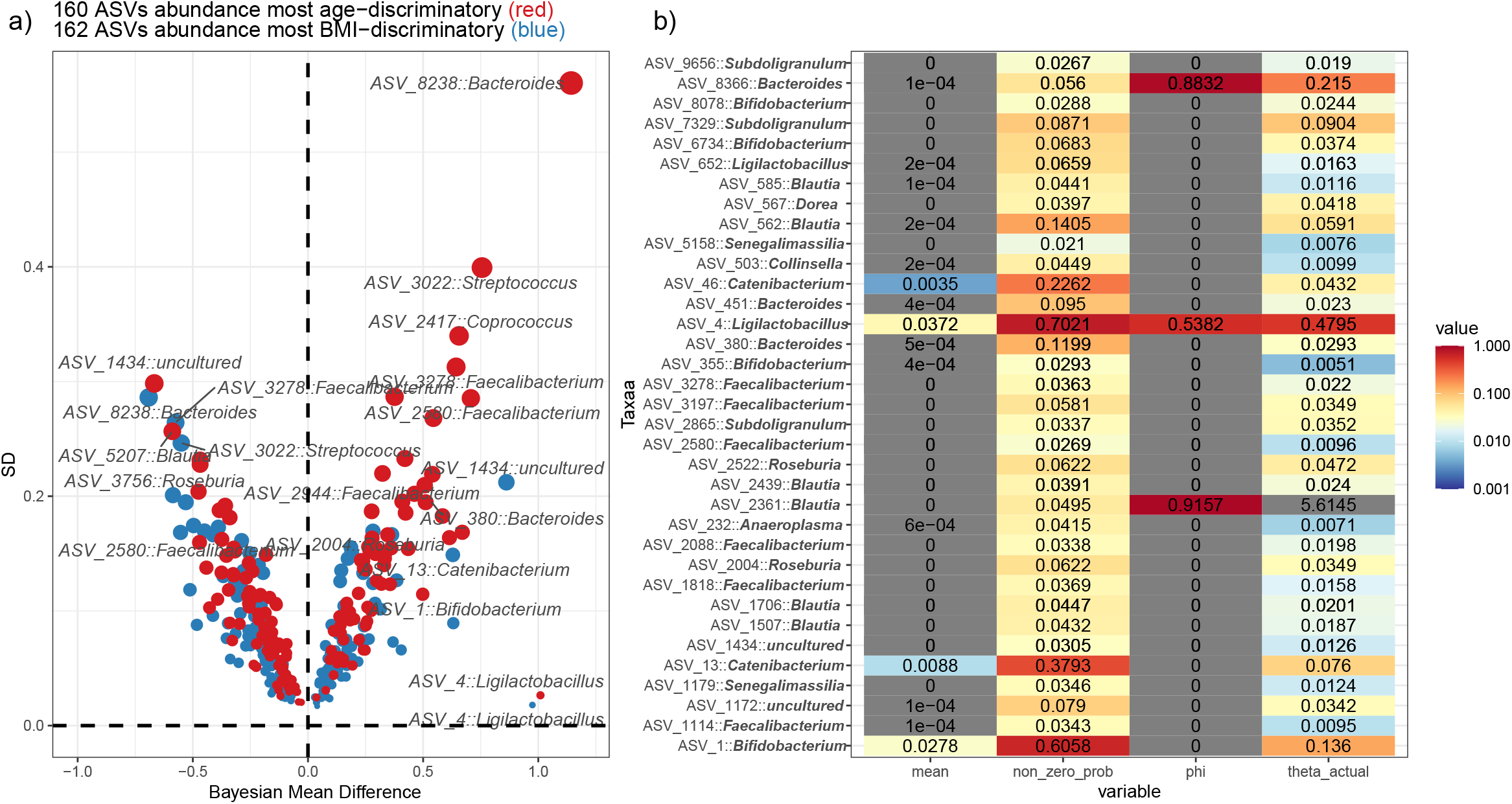
Bayesian effect size analysis was conducted to evaluate the impact of age and BMI on the abundance of taxa. a) The graph plots the influence of age and BMI on taxa, with the effect size represented on the X-axis and the error margin displayed on the Y-axis; Attributes of taxonomic data (ASV), only when its covariate effect sizes of any age or BMI exceeding 0.35, are highlighted, including *ϕ, θ*, average relative abundance, and zero probability.

Therefore, evaluating the ethnic and geographic influence on each ASV allowed us to uncover the diversity and distribution of these microbial species across different populations and regions. Then, we focus on ASVs which covariate effect sizes of any age or BMI exceeding 0.35. These ASVs have at least one pair of non-overlapping ethnogeographic interaction effects (see the Fig. 4). We observed the effects of other ethnic groups by setting the Uyghur ethnic group, located at geographic position H, as the reference control group, in the context of the probability density of the posterior distribution for two covariates, ethnicity and geographic location. We found the discrepancy of interaction between ethnicity and geography in different ASVs. *Ligilactobacillus* (ASV_4) was the most obvious, showing as follows: the positive effect of the Uyghur at location N(effect size : 0.50, 95% CI: [0.20, 0.80]) as well as negative effect of the Kazakh at location N, Uyghur with geographic S and W (effect size : -0.85, 95% CI: [-1.43, -0.26]; -1.53, 95% CI: [-2.58, -0.48]; -0.36, 95% CI: [-0.68, -0.04]). The *Bifidobacterium* (ASV_1) showed similar results. Coincidentally, *Bifidobacterium* and *Ligilactobacillus* genera are the most common consumers of fermented foods and prebiotics [55]. Thus, this prominent association is likely to be explained by regional diets or the heredity of local inhabitants [56]. Indeed, we found that each cohort had their own distinctive eating habits, such as hand-fermented dairy products.

**FIG 4.**
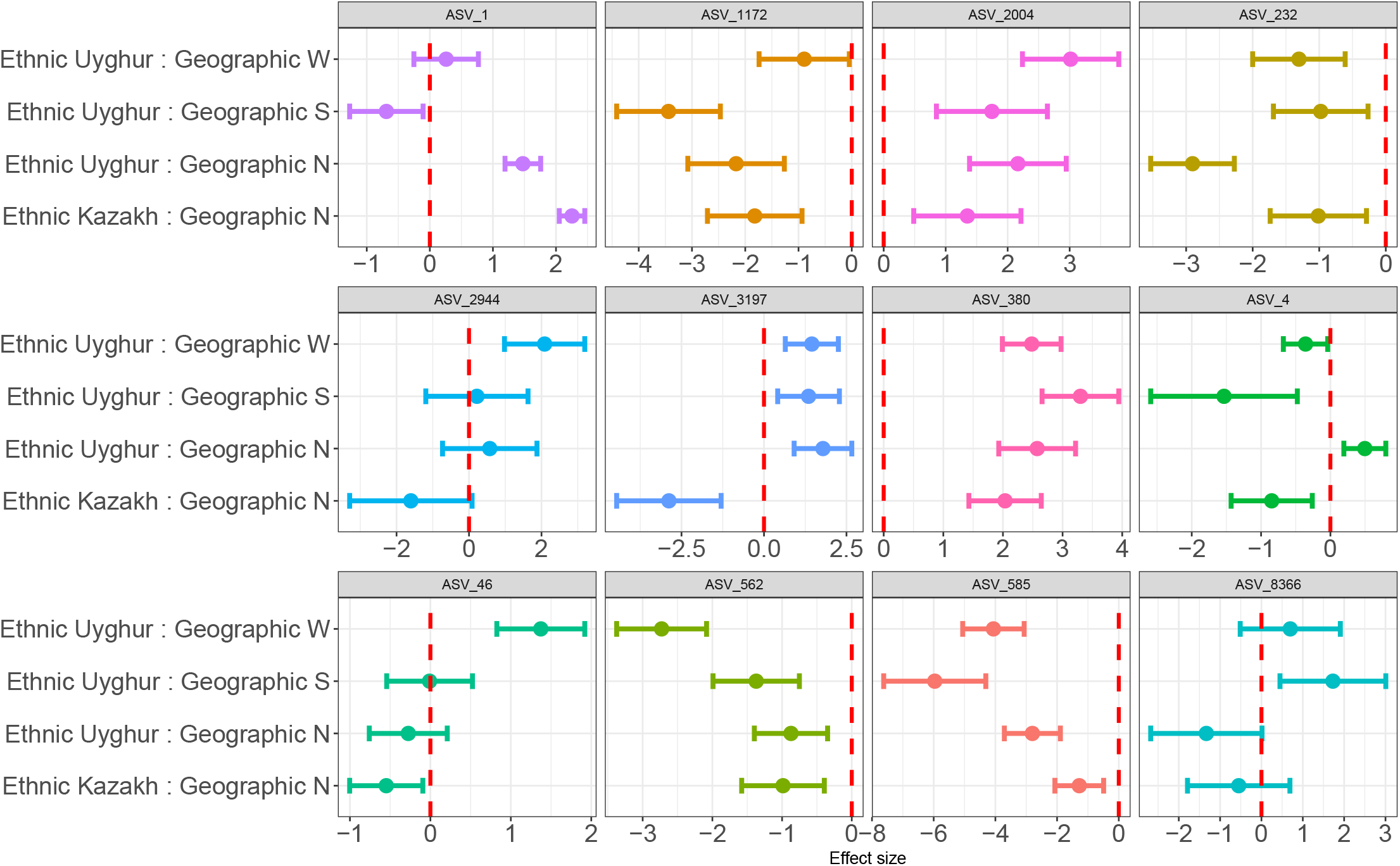
Effect size of 12 taxa under the interaction of ethnicity and geography. Those covariate effect sizes of any age or BMI exceeding 0.35, and at least one pair of ethnogeographic interaction effects do not overlap. The red dotted line represents an effect size of zero.

## Discussion

We proposed and optimized the ZINB model, a method based on Bayesian parametric regression, specifically for modeling counting data in the presence of high zero inflation and high dispersion. We applied this model to studies aiming to dissect gut microbiome count data. The estimates based on the ZINB model provide a more nuanced understanding of how each microbial taxon is affected by covariates, compared to conventional statistical tests alone. The ZINB (*τ*) model has the advantage of being able to accommodate more complex data structures. Zero-inflation is a inevitable property in gut microbiome count data, but previous some models such as BhGLM and glmFit ignore this property and primarily utilize a negative binomial distribution[57]. Conversely, some other studies have used certain statistical models that may not be appropriate, producing results that overemphasize the zero-bloat nature of the gut microbiome data[25, 58, 12, 59]. In the current study, our data processing results by using the ZINB model showed that only a small number of microbial taxa counts belonged to zero-inflated distributions, and the probability of zero-inflation decreases continuously with an increase of taxonomic rank. So, it is important to carefully consider the properties of the data and choose appropriate models that can accurately capture the underlying structure of the data.

In addition to taking the properties of the zero-inflated characteristic into account, ZINB is also capable of considering the degree of dispersion. Compared with previous studies, we found that the degree of dispersion has a significant impact on the accuracy of model parameter estimation, and this effect can be mitigated by increasing the sample size. Our study provides more detailed insights than previous studies into the relationship between dispersion and parameter estimation accuracy. Furthermore, we conducted a detailed discussion on each parameter of ZINB model. We provided insights into the interpretation and practical implications of each parameter in the model. This helps to provide a better understanding of how each parameter affects the model’s performance and how to adjust them to achieve better results in different scenarios. In the initial step of our analysis, we eliminated taxa with fewer than 30 observed counts in the cohort that often lead to model sampling errors. Also, we observe that using the gamma distribution for the parameter *τ* was superior to the Gaussian distribution. Thus, our approach is adaptable since it permits the identification and estimation of the correlation between covariates and the abundance of each taxon. We developed a pipeline for analyzing microbial data using a Zero-Inflated count model was proposed, and applied it to a real cohort study. The pipeline emphasizes strategies for addressing the effect of covariate interactions, such as geography and ethnicity. Compared to adulthood, the microbial composition and body index in childhood tend to undergo relatively dramatic changes. To account for the rapid changes, we performed Z-score normalization on BMI values. Analyzing the variation in BMI values during early life stage requires a more nuanced approach than simply normalizing or centralizing [60] for covariate data.

As for our real-data analysis, we identified differentially abundant taxa in our model that were related to age or obesity, and certain taxa were identified as indicators of interaction with ethnicity or geography, such as the *Bifidobacterium* and *Ligilactobacillus*. Identifying interaction between differentially abundant taxa with ethnicity and geography would help to understand the factors that shape the characteristic human gut microbiome. It is already known that diet plays a decisive role in forming the gut microbiome of people with different genetic backgrounds.Particularly, the consumption of localized fermented foods inhabiting lactic acid bacteria may be related to certain bacterial taxa.

Therefore,these findings have biological significance and help advance our understanding of the mechanisms underlying the gut microbiome. Moreover, developing personalized interventions based on these findings could help promote a healthy gut microbiome in people with diverse backgrounds [61].

Although the zero-inflation negative binomial model is widely acknowledged as the most appropriate way to analyze count-based microbial community profiles, we observed some inconsistent behavior for estimating the probability of extra zeros (Eq 3). This also underscores the potential reproducibility issues that can arise due to variations in algorithms, implementations, and computational environments, even when using the same underlying model [62, 63]. It suggests that interpreting multiple implementations of the same statistical model for complex microbial community settings without an experimentally validated gold standard should be very cautious. Nevertheless, we also observe that the ZINB models suffer from a surplus of random number generation and calculations of myopia: The sensitivity of complex systems to small changes means that model design heavily relies on accurate initial conditions. As reported by Robert McCredie May, the founding father of discrete chaos theory [64]: even the simplest logistic map exhibits an extraordinarily complicated dynamics. Overall, although there are several machine learning techniques recognized for their ability to analyze high-dimensional data and perform feature selection [65, 66, 67], the unique complexity of microbiome count data should not be overlooked when effectively analyzing studies.

Therefore, as nonlinear trajectory-based methods from Bayesian become increasingly available, there is potential for future extensions that address sparse and irregular data, especially When there are multiple covariates [68]. In addition, in microbiome studies, defining an appropriate magnitude for a reasonable and clinically meaningful effect size is unclear. Therefore, there is a need to develop frameworks that make the most of growing amount of microbiome-host interactomics data, facilitating the revelation of underlying biological mechanisms. In summary, the methods presented in our paper provide practitioners with a broad set of effective analytical strategies, with state-of-the-art reasoning capabilities, for identifying microbial associations from complex microbial community in the human microbiome studies.

## Conclusion

In this paper, we propose the ZINB model, which is a Bayesian generalized linear model capable of accounting for multivariate correlation structure, over-dispersion, dimensionality issues, and zero-inflation in the multi-ethnic child gut microbiome data. We detailedly discuss the treatment of covariates, zero-inflation and degree of dispersion in counting data. If these features of gut microbiome data are ignored, it can result in imprecise estimation of effect sizes and reduced statistical power. The results of this study can assist other research groups in making informed decisions when selecting a statistical analysis method.

## ACKNOWLEDGMENTS

This work was supported by the Joint Key Funds of the National Natural Science Foundation of China, and the Autonomous Region Government of Xinjiang, China (Grant No. U1903205), the Science and Technology Special Project of Shihezi Municipal Government (Grant No. 2020PT01), and the Science and Technology Innovation Team Project of Xinjiang Production and Construction Corps (Grant No. 2020CB007). We are grateful to Zhixuan Liang for her instruction in the grammar of the manuscript. J.H. conceived the idea for the manuscript. J.H. and Y.L. are responsible for code editing and figure creation in this study. F.T. and Y.N. are provided valuable guidance on the manuscript. All authors reviewed the manuscript and made corrections.

## DATA AVAILABILITY STATEMENT

The simulation research and pipeline of the Zero-Inflated Negative Binomial (ZINB) model are now available for free on https://github.com/jianhuang525/ZINBmodel.

## ETHICS APPROVAL

This study involves human participants and the study was approved by the Ethics Committee of the First Affiliater Hospital, Shihezi University School of Medicine (2017-117-01). Participants gave informed consent to participate in the study before taking part.

## CONFLICTS OF INTEREST

The authors declare no conflict of interest.

### Supplemental material

#### S1 Appendix

**Supplementary algorithms. estimate zero-inflated probability and** *ϕ* **parameter**. For the bacteria count data, we estimate the data properties of each ASV/OTU to determine whether it conforms to the zero-infa negative binomial distribution. It is vitally important for us to choose the right model. According to the Formula (1) of the text part, the variance and mean are knowable. At this point, the mean and variance in the negative binomial distribution are:

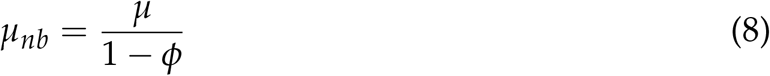

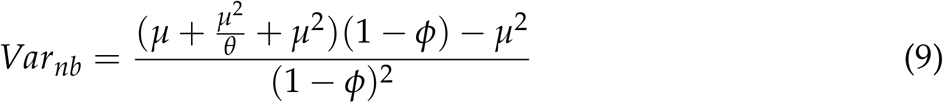

By utilizing the variance and mean, we can determine the dispersion parameters *θ* of y.

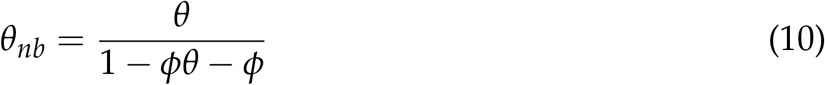

In this way, we can calculate the true probability of zero expansion:

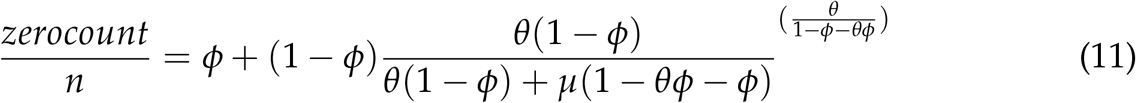

According to the above equation, we can solve the *ϕ* parameter.

**Table S1 Child cohort information**. The table presents information on 585 children, including sample number, geographic location, ethnicity, age, gender, and BMI for age z score.

**Table S2 Child cohort ASV table**. ASV form data sharing, easy to repeat data processing parts.

**Table S3 Species taxonomy table**. Taxonomy table, easy to repeat data processing parts.

**Table S4 Estimate of the Bayesian effect size**. Shown that 160 ASVs abundance remain age-discriminatory and 162 ASVs abundance remain BMI-discriminatory. And the taxonomy of these ASVs at the generic level.

